# Robust nonparametric descriptors for clustering quantification in single-molecule localization microscopy

**DOI:** 10.1101/071381

**Authors:** Shenghang Jiang, Sai Divya Challapalli, Yong Wang

## Abstract

We report a robust nonparametric descriptor, *J*′(*r*), for quantifying the spatial organization of molecules in singlemolecule localization microscopy. *J*′(*r*), based on nearest neighbor distribution functions, does not require any parameter as an input for analyzing point patterns. We show that *J*′(*r*) displays a valley shape in the presence of clusters of molecules, and the characteristics of the valley reliably report the clustering features in the data. More importantly, the position of the *J*′(*r*) valley (*r*_*J*′_*m*__) depends exclusively on the density of clustering molecules (*ρ*_c_). Therefore, it is ideal for direct measurements of clustering density of molecules in single-molecule localization microscopy.

## Introduction

Single-molecule localization microscopy (SMLM) has been utilized broadly in imaging biological molecules – proteins, DNA, and RNA – in various biological systems.^1–5^ More importantly, by localizing individual molecules, SMLM has allowed quantitative analyses on the spatial organizations and patterns of these molecules, and produced new, quantitative and crucial information that was not accessible previously. New mechanisms of various cellular and molecular organizations and activities at the single-cell level have been unraveled using SMLM.^6–15^

Many algorithms have been adopted, utilized, or developed, in the field of SMLM for analyzing localization data of molecules and quantifying inter-molecular organizations.^13, 14, 16–23^ These methods provide means to distinguish single molecules from clusters of molecules, to examine complex patterns of molecular organization, and to quantify features of spatial organizations. For example, pair-correlation analysis has been applied to SMLM data on membrane proteins to detect the presence of clusters, as well as various cluster features, such as the density of molecules in a cluster and overall size of a cluster.^16^ In addition, density-based algorithms such as DBSCAN (density-based spatial clustering of applications with noise)^24, 25^ and OPTICS (ordering points to identify the clustering structure)^26, 27^ have been exploited to identify clusters of proteins and nucleic acids, as well as to probe the clustering structures, in both bacteria and animal cells.^13, 14, 17–19^ Another method that has been used for analyzing SMLM data is Ripley's K function and its derivatives.^20, 21^ More recently, Bayesian analysis and Voronoï diagrams have been utilized to identify and analyze the clustering of biological molecules.^22, 23^

However, a general problem in some of these methods or algorithms is the need of input parameters. For example, DBSCAN and OPTICS need two parameters (a radius, eps, and the minimum number of points in the neighborhood for a point to be considered as a core point, minPts),^24–27^ and they are known to be sensitive to the chosen parameters.^18, 28^ The identification of clusters in the Voronoï diagram based method also requires a density threshold to determine whether points form clusters.^23^ Although various techniques have been proposed to determine “appropriate’’ parameters for use,^23, 25, 27, 29^ bias is inevitably introduced by the choice of parameters in these algorithms. As a result, a robust nonparametric method is desired for the quantification of clustering features of molecules.

Here we present a descriptor based on nearest neighbor distribution functions for quantifying the spatial pattern of molecules in SMLM. We examined nearest neighbor function,^30^ *G*(*r*), spherical contact distribution function,^30^ *F*(*r*), and the J-function,^31, 32^ *J*(*r*) = (1−*G*(*r*))/(1-*F*(*r*)), and found that the associated derivative functions, *G*′(*r*) and *J*′(*r*), reliably report the clustering features of points. In the presence of clusters, *G*′(*r*) and *J*′(*r*) are peak/valley shaped. More importantly, we observed that 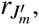 the position of the *J*′(*r*) valley, depends exclusively on the density of clustering points (*ρ*_c_). Therefore, it is ideal for direct measurements of the clustering density of molecules in SMLM. We expect that this nonparametric descriptor *J*′(*r*) is useful in a broad range of applications in SMLM.

## Results

### *G*(*r*), *F*(*r*) and *J*(*r*), and their derivatives

When quantifying the spatial organization of biological molecules in SMLM data, of particular interest in certain situations is the clustering or aggregation of molecules,^33–36^ which is featured by an enhancement in the local density of molecules. This enhancement in density has been used to identify clusters methods such as DBSCAN, OPTICS, and Voronoï tessellation.^13, 14, 17–23^ On the other hand, the enhancement in the molecular density is also accompanied by the decrease of intermolecular distances, which could be described by functions based on nearest neighbor distances, such as pair-wise correlation function,^16^ nearest neighbor function G(r), and spherical contact distribution function *F*(*r*).^30^ The nearest neighbor function *G*(*r*) is the distribution function of the distance r of a point (existing in the data) to the nearest other point, while the spherical contact distribution *F*(*r*) is the distribution function of the distance r of an arbitrary point in the space (not necessarily existing in the data) to the nearest point in the data.^30^ In addition, another function, *J*(*r*), has been suggested by van Lieshout and Baddeley in 1996,^31^ 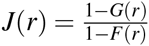, as a better nonparametric test to determine whether data were from a Poisson process.

We first explored how *G*(*r*), *F*(*r*) and *J*(*r*) functions depends on the clustering features of points using numerical simulations. Briefly, we generated points forming various clusters in the presence of noises (i.e., Poisson random points) in a region of interest, and computed these three functions. In a two-dimensional Poisson random process where points were not forming clusters (Fig. 1A), the nearest neighbor functions gave the expected curves, 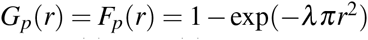 and *J*_*p*_(*r*) = 1 (Fig. 1C). However, when points aggregated into clusters (Fig. 1B), both *G*(*r*) and *J*(*r*) deviated significantly from random points, while *F*(*r*) became only slightly different (Fig. 1D). We observed that *J*(*r*) drops quickly from 1 to ∼0.4 when *r* increases from 0 to 5 nm, while *G*(*r*) raises quickly in the same r-range (0–5 nm). This observation indicates that *G*(*r*) and *J*(*r*) could be used for detection of clusters.

**Figure 1.**
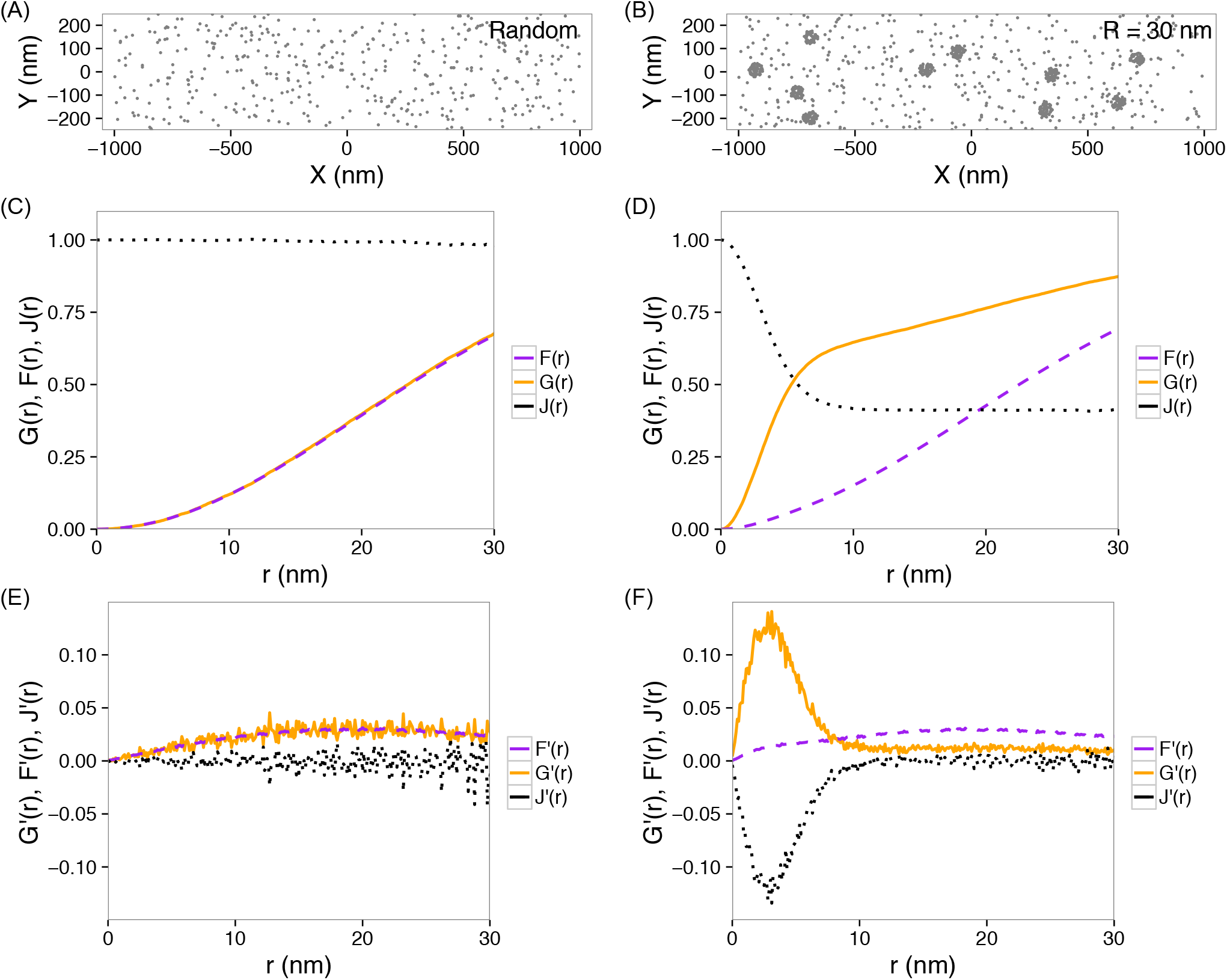
*G*(*r*), *F*(*r*) and *J*(*r*) functions, and their derivatives. (A) Simulated noise points. (B) Simulated points forming clusters with a radius of *R* = 30 nm, in the presence of noise points. (C, D) *G*(*r*), *F*(*r*) and *J*(*r*) functions calculated from the points in (A) and (B), respectively. (E, F) Derivatives, *G*′(*r*), *F*′(*r*) and *J*′(*r*), calculated from the points in (A) and (B), respectively.

Furthermore, to remove accumulative effects, and inspired by Kiskowski et.al.,^37^ we calculated the derivatives of these functions (*G*′(*r*), *F*′(*r*) and *J*′(*r*)). Striking peaks or valleys appeared in *G*′(*r*) and *J*′(*r*) if points formed clusters (Fig. 1F). In contrast, these derivative functions remain essentially flat for random points (Fig. 1E). On the other hand, *F*′(*r*)’s were very similar in the two cases (Fig. 1E–F).

### Dependence of *G*′(*r*) and *J*′(*r*) functions on clustering features

To explore quantitative applications of *G*′(*r*) and *J*′(*r*), we examined how they differ when the clustering features of points vary. Here we focus on the following features: the radius of clusters, *R*_*c*_, the density of points in clusters, *ρ*_*c*_, the number of clusters, *n*_*c*_, the density of (random) noise points, *ρ*_*r*_, and the width (*W*) and height (*H*) of the region of interest. The first three features, *R*_*c*_, *ρ*_*c*_ and *n*_*c*_, are directly related to the properties of clusters in the data, while *ρ*_*r*_ is an indicator of the noise level. By varying one feature at a time, we observed that changes in *ρ*_*c*_, *ρ*_*r*_, or *R*_*c*_ resulted in horizontal shifting or vertical scaling of both *G*′(*r*) and *J*′(*r*) (Fig. 2A–C). For example, both *G*′(*r*) and *J*′(*r*) shift to the left and scale up when the clusters become denser (*ρ*_*c*_ increases). If the clusters become bigger (*R*_*c*_ increases) while keeping the clustering density constant, little horizontal translation was observed (Fig. 2C), although both *G*′(*r*) and *J*′(*r*) scale up too. In contrast, *G*′(*r*) and *J*′(*r*) are insensitive to the number of clusters (*N*_*c*_) or the size of the region of interest, *W* and *H* (Fig. 2D–F).

**Figure 2.**
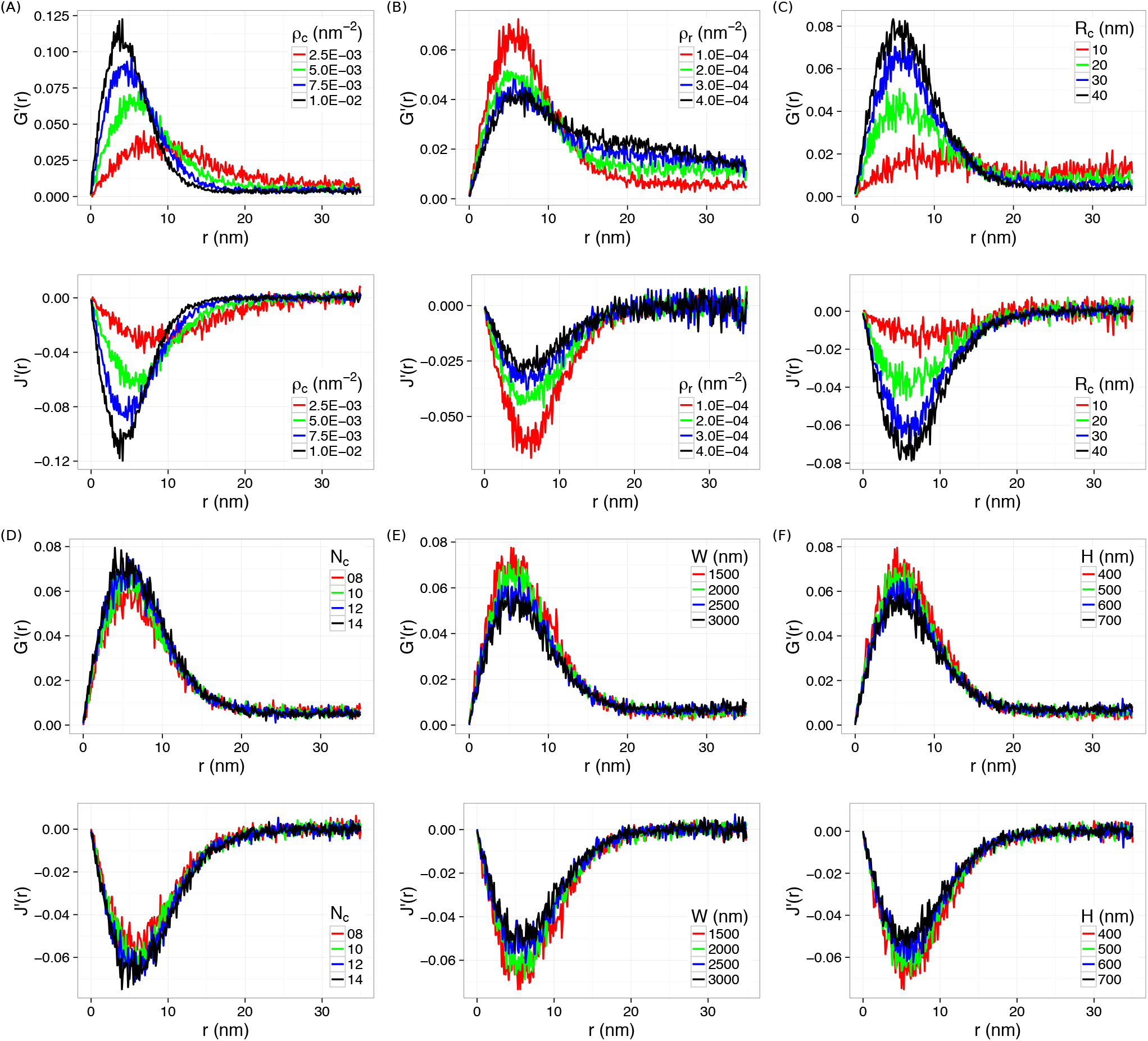
Changes in *G*′(*r*) and *J*′(*r*) by varying a cluster feature at a time. (A) *ρ*_*c*_, (B) *ρ*_*r*_, (C) *R*_*c*_, (D) *N*_*c*_, (E) *W*, and (F) *H*.

We further quantified the dependence of *G*′(*r*) and *J*′(*r*) on the clustering features. By fitting *G*′(*r*) and *J*′(*r*) with polynomials, both the amplitude (height *G*′_*m*_ for *G*′(*r*), or depth *J*′_*m*_ for *J*′(*r*)) and the positions (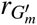 and 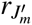) of the peaks and valleys were determined. The dependence of these values on the clustering features are shown in Fig. 3, S1, S2 and S3. We observed that both *G*′_*m*_ and *J*′_*m*_ depend on all the clustering features (Fig. S1 and S3), but 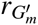 and 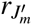 are most sensitive to the density of clustering points *ρ*_*c*_ (Fig. 3 and S2). Most interestingly, 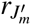 is essentially independent on all the clustering features except the density of clustering points *ρ*_*c*_ (Fig. 3), providing a way to correlate 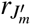 with directly measuring the clustering densities of molecules, as shown below.

**Figure 3.**
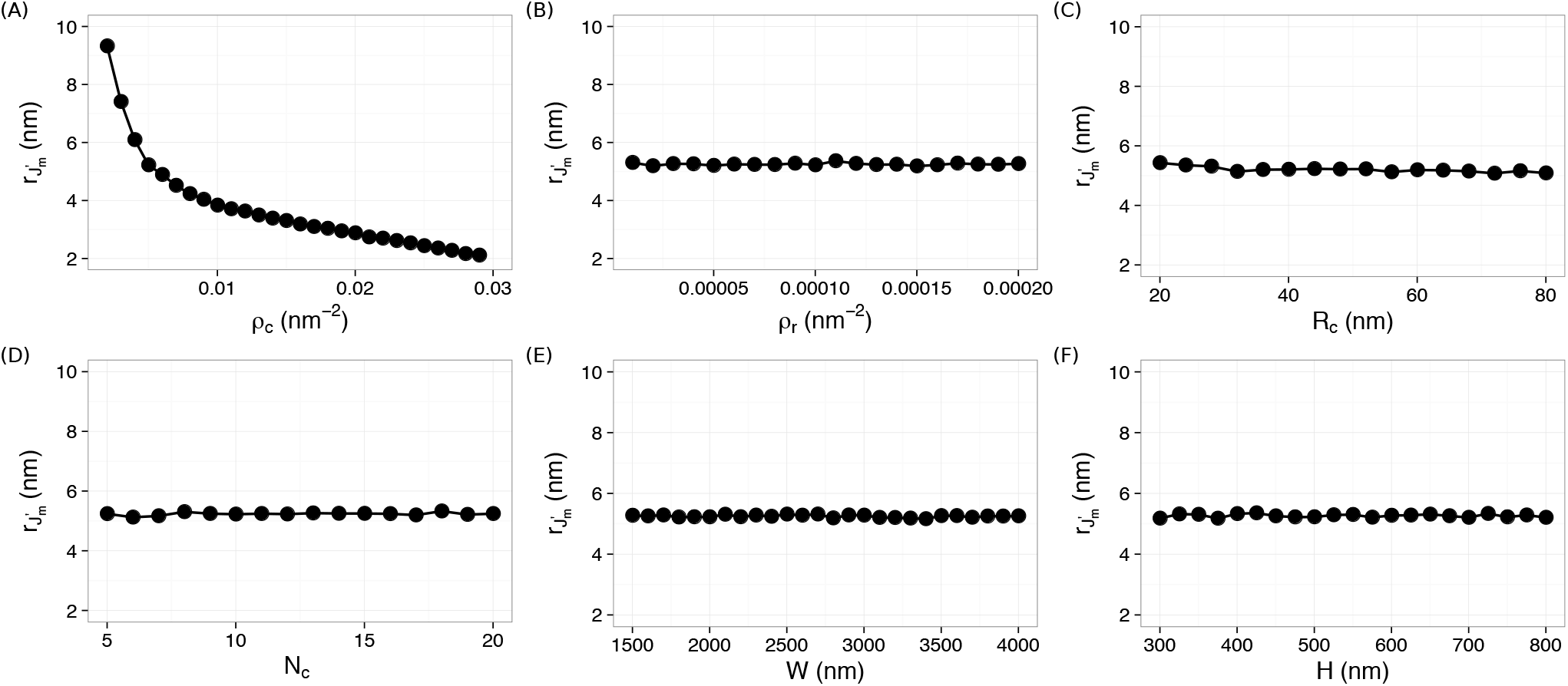
Dependence of 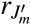 on the clustering features: (A) *ρ*_*c*_, (B) *ρ_r_*, (C) *R*_*c*_, (D) *N*_*c*_, (E) *W*, and (F) *H*.

### Direct measurement of the density of clustering points by 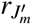

To explore the possibility of using 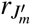 for direct measurements of clustering densities of molecules, we first confirm that the 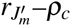 relation is independent on other clustering features even if we simultaneously vary both *ρ*_*c*_ and *R*_*c*_, or *N*_*c*_, or *ρ*_*r*_…. We found that the the 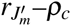 relation from all the simulations collapse onto a single curve (Fig. 4A). Fitting all the data with a power-law function 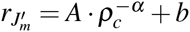 gives α = 0.76±0.03, yielding a “calibration’’ curve for measuring clustering densities in any point patterns. We note that the 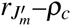 relation is monotonic over a larger range of *ρ*_*c*_ (Fig. 3A); however, more complicated mathematical forms other than a single power-law are needed to fit the “calibration” curve.

**Figure 4.**
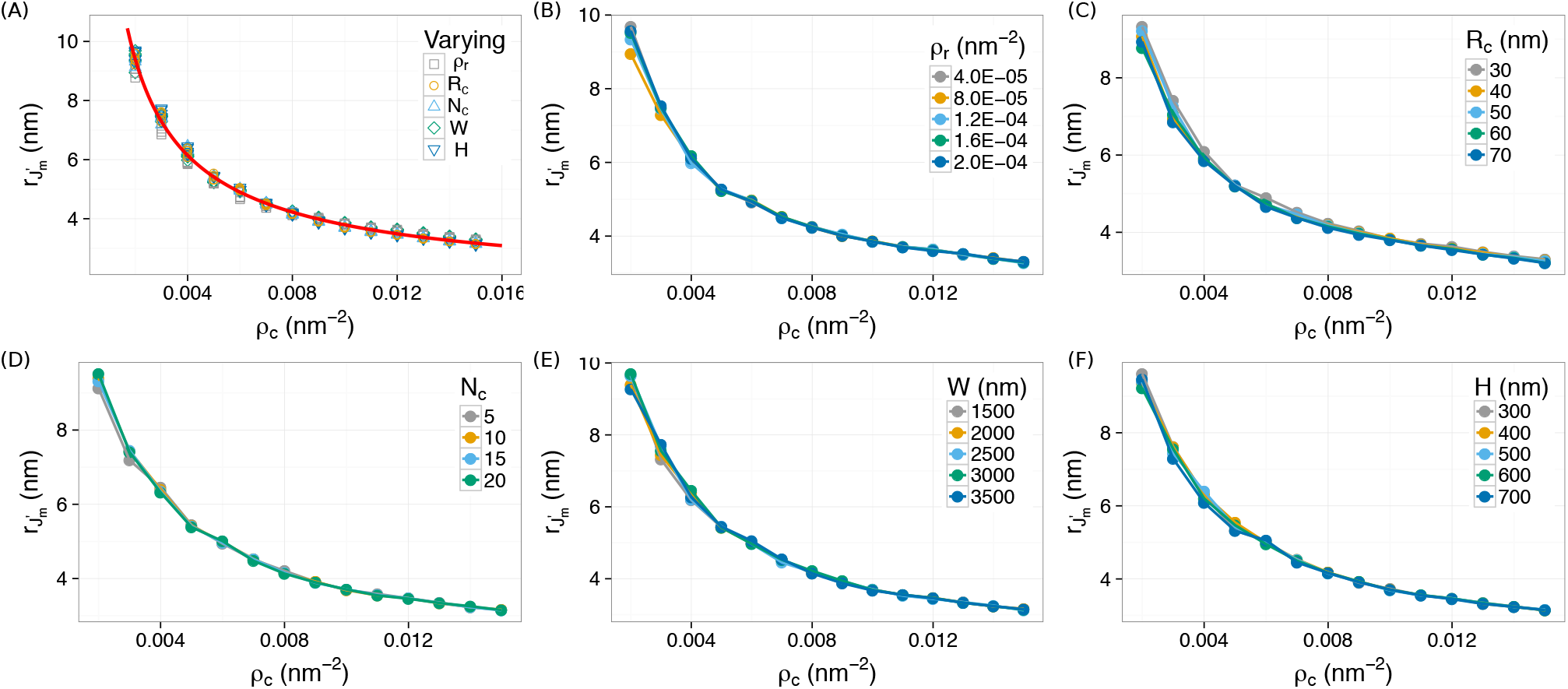
The 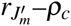 relation is independent on all the other cluster features, *R*_*c*_, *ρ*_*r*_, *N*_*c*_, *W*, and *H*. All data points collapse onto a single power-law curve, 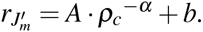 Least-square fitting gives 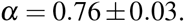

To verify the capability of using the 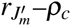 “calibration’’ curve to directly measure the clustering density of molecules in SMLM data, we exploited numerical simulations again, for which we know the ground-truth clustering density. Briefly, we generated 50 simulated data with a “unknown” clustering density (*ρ*^*t*^_*c*_ = 0.0103 nm^™2^), and computed *J*′(r) and 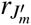 for each simulation. The “measured” clustering density 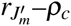 is then obtained from the 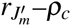 “calibration” curve, 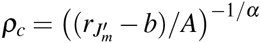. We found that the “measured” clustering densities 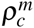 are very close to the the ground-truth density 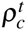: the relative error, 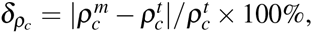 is below 4.6% for all the 50 simulations, and the average relative error is 2.5%.

### Robustness of the 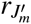 relation

“Noises” are almost always present in SMLM data, due to individual molecules not forming clusters, non-specific labeling, and/or false-positive localizations. Amazingly, we found that 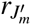 is independent on the density of random points in the data, as shown in Fig. 3B and 4, strongly suggesting that the use of 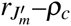 relation for measuring clustering densities is robust.

To further assess the robustness of the 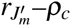 relation, we systematically investigated, given a density of clustering point (*ρ*_*c*_), how 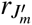 deviates in the presence of various amount of noises. The noise level is defined as *β* = *n*_*rp/*_ *n*_*cp*_, where *n*_*rp*_ is the number of (random) noise points, and *n*_*cp*_ is the number of clustering points, in the whole region of interest. We found that *rJ*′_*m*_ remains constant (with relative errors 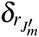 below 5%) even if there are ≈10 times more noise points than clustering points (Fig. 5), demonstrating that the 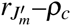 relation is extremely robust.

**Figure 5.**
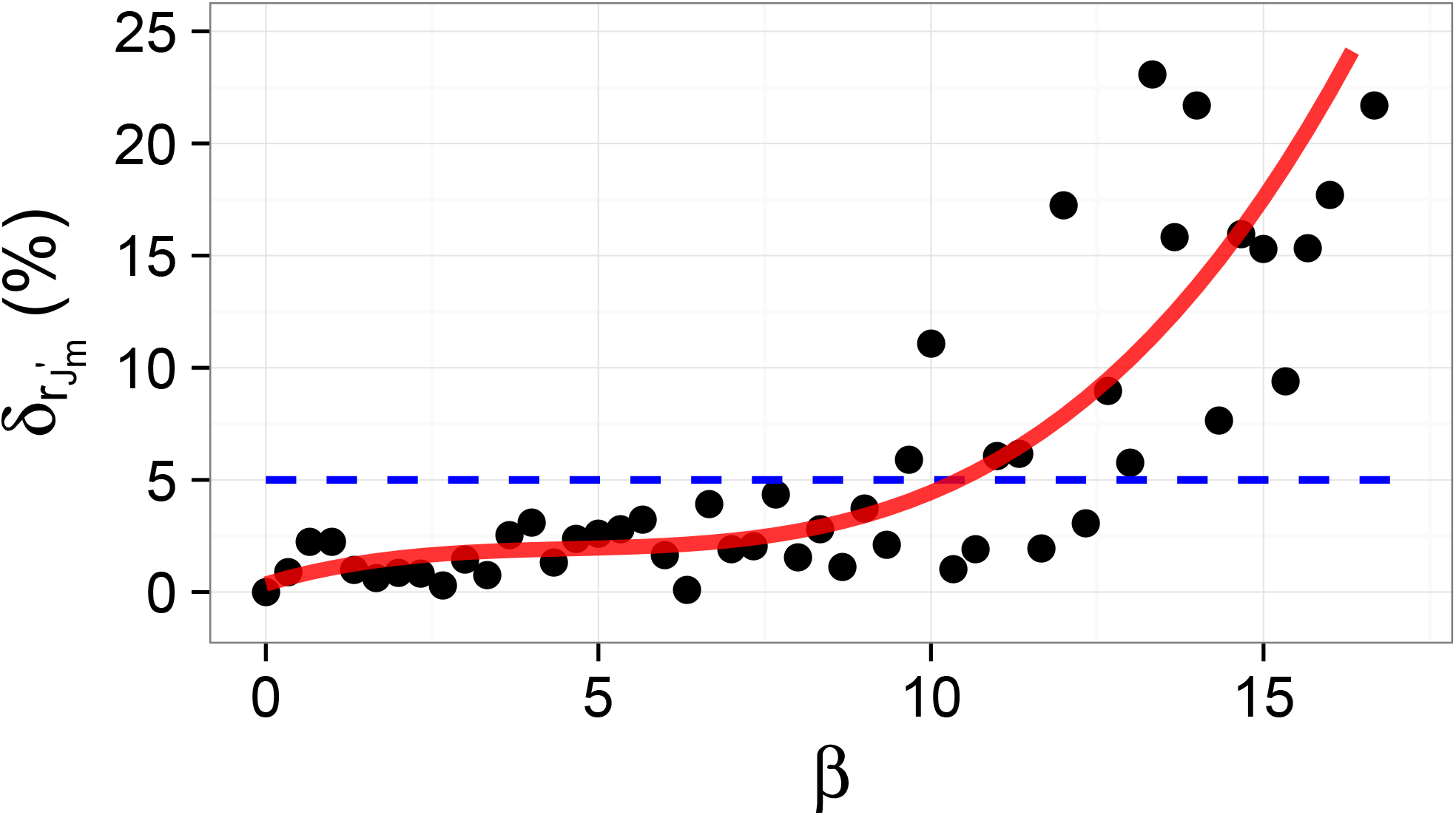
Robustness of the 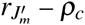 relation. Given a certain density of clustering points (*ρ*_*c*_), 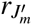 remain (almost) constant (i.e., the relative error 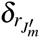 is < 5%) until there are 10 times more noise points than clustering points (*β* ≤ 10).

### *J*′(*r*) as a global descriptor for heterogeneous clusters

It's known that, in certain applications, molecules of interest might form heterogeneous clusters.^16, 23^ We examined heterogeneity arising from either clustering radius (*R*_*c*_) or clustering density (*ρ*_*c*_). Briefly, simulations were run for clusters with two different clustering radii (*R*_*c*1_ and *R*_*c*2_), or two different clustering densities (*ρ*_*c*1_ and *ρ*_*c*2_), in the presence of random noises. We noticed that 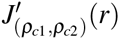 from heterogeneous clusters with different clustering densities shifted both horizontally and vertically, and fell between the two curves from homogeneous clusters, 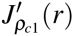 and 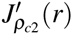 (Fig. 6). In addition, we observed that 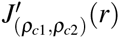 overlapped very well with 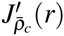 from a homogeneous sample with a clustering density equal to the algebraic mean, 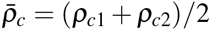 (Fig. 6). It is noted that 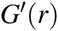 shows a similar behavior. In contrast, for heterogeneous clusters with different radii, 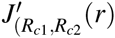 shifted only in the vertical direction. 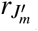 does not change for heterogeneous clusters with different radii (Fig. S4), which is expected because the 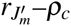 relation does not depend on *R*_*c*_ anyway. In addition, we found that 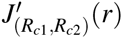 is equivalent to 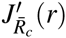 from homogeneous clusters with a radius of 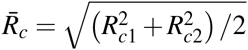 (Fig. S4). In summary, we conclude that *G*′(*r*) and *J*′(*r*) are descriptors reporting the “global” clustering density throughout the region of interest.

**Figure 6.**
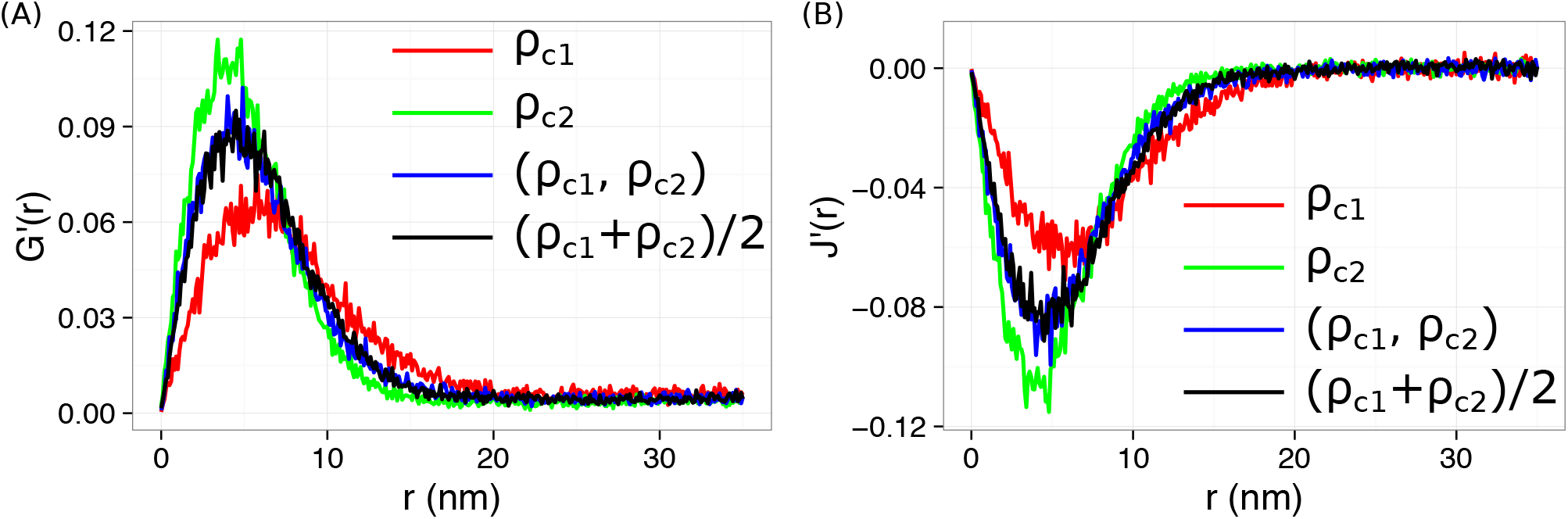
*G*′(*r*) and *J*′(*r*) for data with heterogeneous clusters with two different clustering densities.

## Discussion

To conclude, we explored the possibility of utilizing nearest neighbor functions to quantify spatial patterns of molecules in single-molecule localization microscopy. We observed that the associated derivative functions, *G*′(*r*) and *J*′(*r*), could reliably report the clustering features of point patterns. We found that *J*′(*r*) is particularly useful because its position, 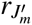, relies exclusively on the density of clustering points (*ρ*_*c*_). More importantly, we showed that this 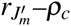 relation is very robust in the presence of up to∼10 times more noise points than clustering points.

In the current study, we chose not to exploit any border correction when computing the nearest neighbor functions. A simplest approach for border correction is the “reduced sample” method,^38^ which focuses on the points lying more than *r* away from the boundary of the region of interest. However, the “reduced sample” method discards much of the data, and therefore unacceptably wasteful. In addition, it's particularly inappropriate in certain applications where points are preferentially located at the boundary, an example of which is the spatial organization of high-copy number plasmids in bacteria.^14^ We note that more sophisticated methods for border correction are available, including the Kaplan-Meier correction^39^ and the Hanisch correction,^40^ both are provided in the *spatstat* R-package.^41, 42^ These edge corrections can readily used in our method. However, for the sake of simplicity, uncorrected estimators for the nearest neighbor functions have been used in the current study.

We would like to emphasize that the current method based on nearest neighbor functions is nonparametric and robust. Computing the nearest neighbor functions and their derivatives does not require any parameters as human inputs, eliminating possible subjective biases that might exist in other algorithms such as DBSCAN and OPTICS. In addition, the performance of this method is robust, even in the presence of 10 times more noise points than clustering points. The nonparametric nature and robustness of the current method would allow broad applications in the field of single-molecule localization microscopy.

We expect several types of applications of our method in the field of SMLM. First, it can be used as a direct quantification of the clustering density (*ρ*_*c*_) of molecules in biological samples. Second, although our method does not identify clusters by itself, it provides objective means to determine parameters (i.e., clustering density) that could be used in other clustering-identification algorithm such as DBSCAN and Voronoï tessellation. In addition, in the current work, we focused on the 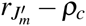 relation for non-parametric measurement of the clustering density of molecules; however, it's possible to design a way to figure out other cluster features (such as *R*_*c*_ and *ρ*_*r*_) by taking advantage of the dependence of *G*′_*m*_ and *J*′_*m*_ on those features (Fig. S1 and S3), together with the information of *ρ*_*c*_.

Finally, we would like to mention that, although the current study shows that the nearest neighbor functions are global descriptors, which cannot dinstinguish heterogeneous clusters, more sophisticated algorithms based on the nearest neighbor functions might be developed in order to detect heterogeneities in the data. For example, it's possible to divide the region of interest into sub-regions, perform computations on *J*′(*r*), and report heterogeneous features on different sub-regions.

## Methods

### Spherical contact distribution function *F*(*r*), nearest-neighbor distribution function *G*(*r*), and the *J* function *J*(*r*)

In a set of points, *X*, in the *k*-dimensional space, the spherical contact distribution function, or sometimes referred to as the empty space function, *F*(*r*), of *X* is defined as 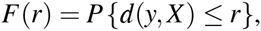, where 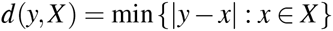 is the distance from an arbitrary point, *y*, to the nearest point of the point process, *X*.^30^ For a Poisson process in the *k*-d space, 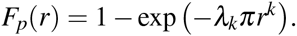.^30^ The nearest-neighbor distribution function *G*(*r*) is very similar to 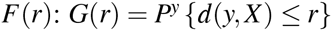 where *P*^*y*^ is the Palm distribution, which is the conditional distribution of the entire process given that *y* is one point in *X*.^30^ Therefore, *G*(*r*) is the distribution function of the distance from a point of the process to the nearest other point of the process, i.e., the “nearest-neighbor”. For a Poisson process in the *k*-d space, 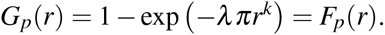.^30^ In 1996, van Lieshout and Baddeley suggested using the quotient 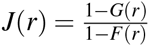 to characterize a point process.^31^ For a Poisson process, 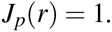.

### Simulation and computation of *G*(*r*), *F*(*r*), *J*(*r*) and their derivatives

Sets of points were generated in R programing language.^43^ In a region of interest with a width (*W*) and a height (*H*), *n*_*c*_ circular clusters with radii of *R*_*c*_ were randomly distributed. Each cluster contains random points at a density of *ρ*_*c*_. Poisson noise points were added randomly to the whole region of interest, with a density *ρ*_*r*_. The total number of clustering points 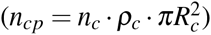 and the total number of noise points 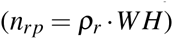 define the noise level *β* = *n*_*rp*_*/n*_*cp*_.

Simulations were run using various sets of cluster features (*W*, *H*, *ρ*_*r*_, *ρ*_*C*_, *n*_*c*_, *R*_*c*_). For each set of features, 50–200 trials were run. The *G*(*r*), *F*(*r*), *J*(*r*) functions and their derivatives were computed using the *spatstat* package,^41, 42^ without applying any edge corrections.

## Acknowledgments

This work was supported by the Human Frontier Science Program (LT000752/2014-C to Y.W.) and by the Arkansas Biosciences Institute, the major research component of the Arkansas Tobacco Settlement Proceeds Act of 2000.

## Author contributions statement

Y.W. conceived the project, S.J. and Y.W. conducted the simulations, S.J., S.D.C. and Y.W. analyzed the results. S.J., S.D.C. and Y.W. wrote and reviewed the manuscript.

## Additional Information

### Corresponding author

Correspondence to Yong Wang.

## Supplementary Information

**Figure S1.**
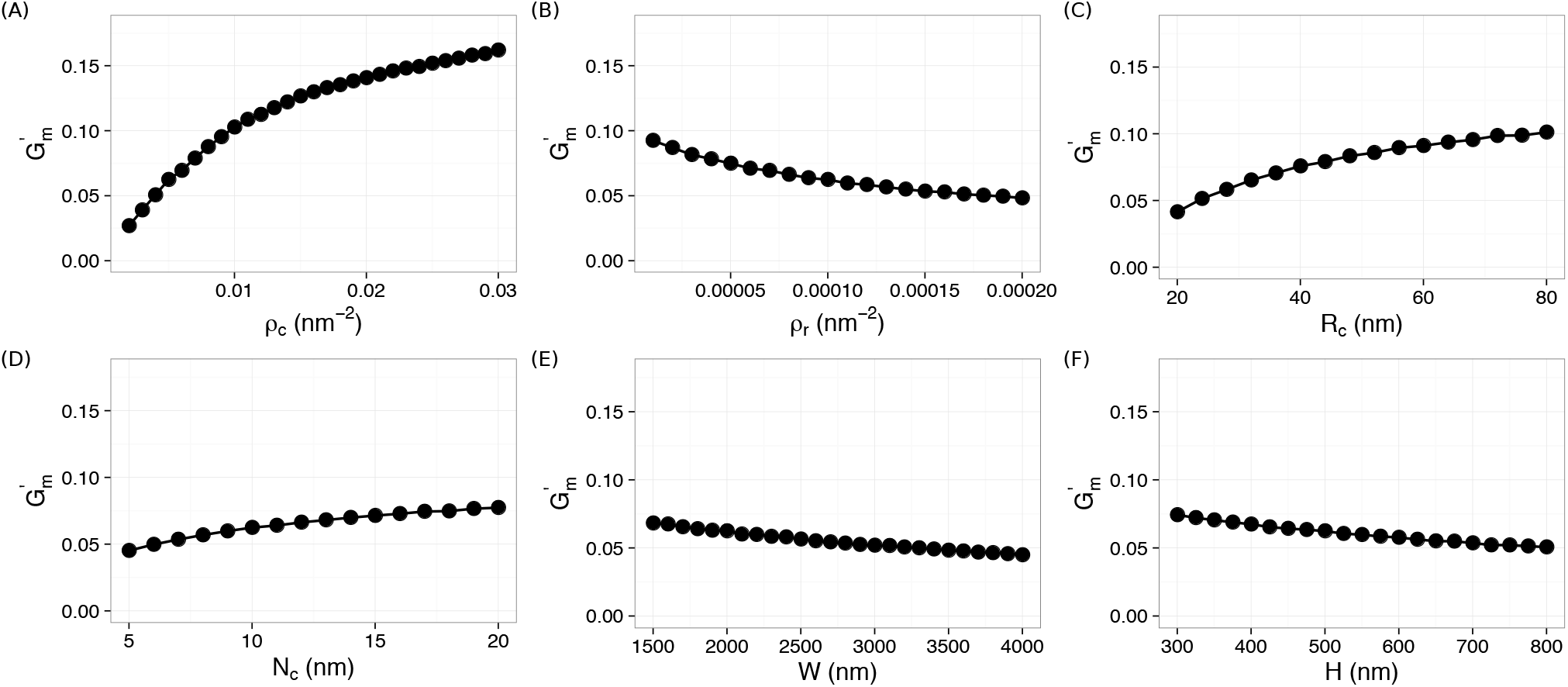
Dependence of 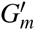 on the clustering features: (A) *ρ*_*c*_, (B) *ρ*_*r*_, (C) *R*_*c*_, (D) *N*_*c*_, (E) *W*, and (F) *H*.

**Figure S2.**
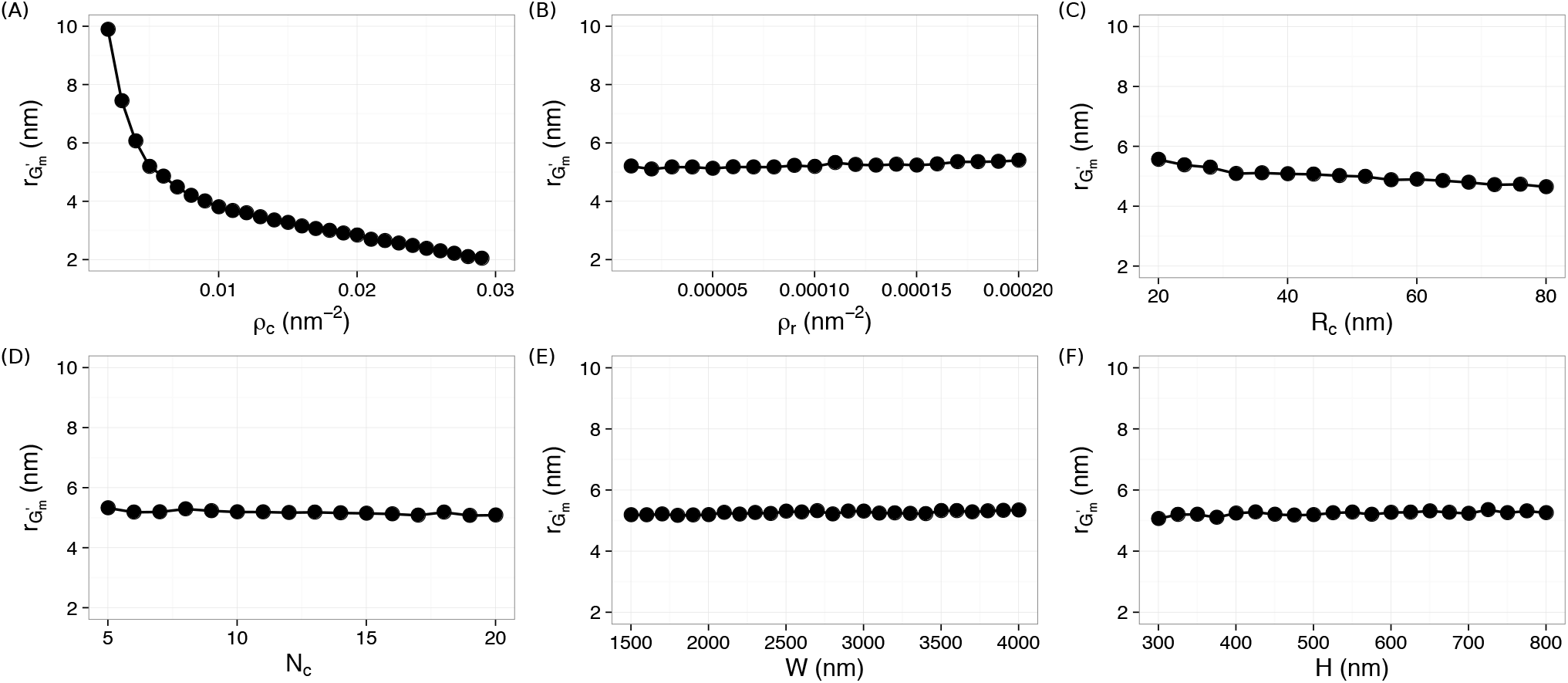
Dependence of 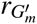 on the clustering features: (A) *ρ*_*c*_, (B) *ρ*_*r*_, (C) *R*_*c*_, (D) *N*_*c*_, (E) *W*, and (F) *H*.

**Figure S3.**
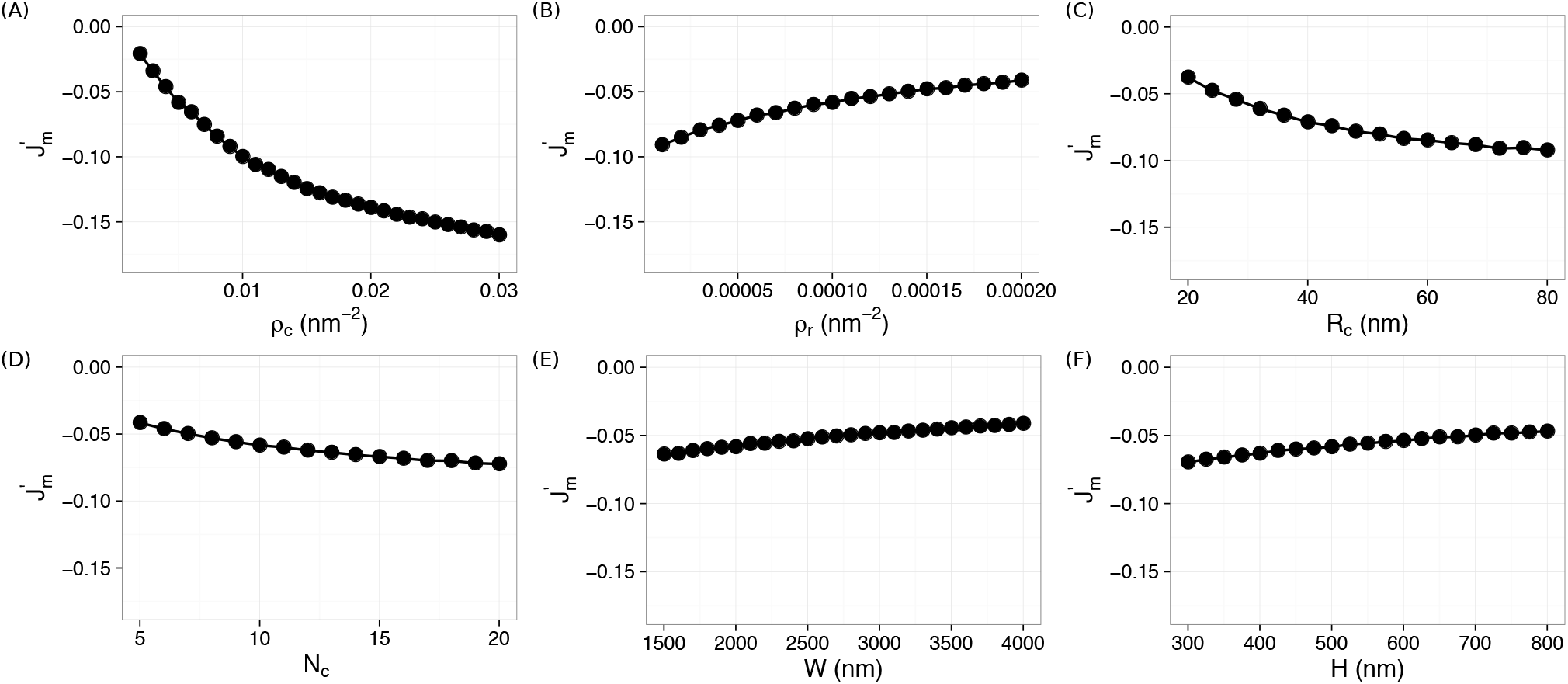
Dependence of 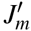 on the clustering features: (A) *ρ*_*c*_, (B) *ρ*_*r*_, (C) *R*_*c*_, (D) *N*_*c*_, (E) *W*, and (F) *H*.

**Figure S4.**
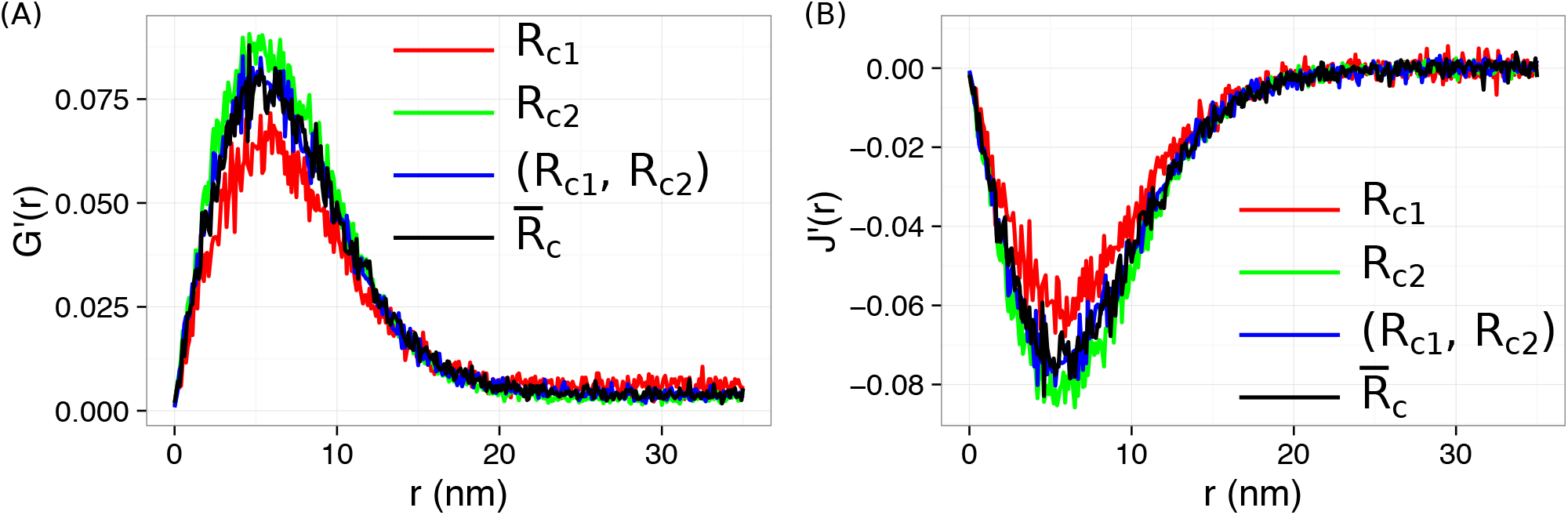
*G*′(*r*) and *J*′(*r*) from heterogeneous clusters with different radii, 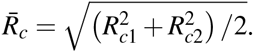.

